# Link between personality and response to THC exposure

**DOI:** 10.1101/674044

**Authors:** Tetiana Kardash, Dmitry Rodin, Michael Kirby, Noa Davis, Igor Koman, Jonathan Gorelick, Izhak Michaelevski, Albert Pinhasov

**Author notes:** Corresponding Author: Prof. Albert Pinhasov, Department of Molecular Biology, Ariel University, Ariel, Israel 4070000. **Abbreviations:** Dom, dominant; Sub, submissive; THC, delta-9-tetrahydrocannabinol; CPP, condition place preference; RS, Restraint Stress; TST, Tail Suspension Test; FST, Forced Swim Test; DSR, Dominan Submissive Relationship; CORT, corticosterone; HPA, hypothalamic-pituitary-adrenal axis.

## Abstract

The effects of cannabis reported by users range from experiences of euphoria and anxiolytic effects to paranoia, anxiety, and increased risk of depression. Attempts to reconcile the apparent contradictions in user response have not been conclusive. Here, we utilized selectively-bred stress-resilient socially dominant (Dom) and stress-sensitive socially submissive (Sub) mice to elucidate this contradiction. Following short-term, repeated treatment with delta-9-tetrahydrocannabinol (THC) at two different doses (1.5 mg/kg and 15 mg/kg), Sub mice presented significant place-aversion in a Conditioned Place Preference paradigm at a high dose, whereas Dom mice displayed no place preference or aversion. Forced Swim test conducted after 6-week of washout period, revealed differential impact of the two THC doses depending upon behavioral pattern. Specifically, the low dose alleviated depressive-like behavior in Sub mice, while the high dose produced the opposite effect in Dom mice. Interestingly, corticosterone concentration in serum was elevated at the high dose regardless of the mice-population tested. We conclude here that differences in dominance behavior and stress vulnerability are involved in the regulation of cannabis response among users and should be considered when prescribing THC-containing medications to patients.

## 1. Introduction

Cannabis is the most widely-used drug worldwide with an estimated 183 million users, which equates to 3.8% of adults aged 15–64 years [1]. Several factors have contributed recently to an improved epidemiological profile of cannabis use. Increase of cannabis use for medical purposes [2-8], higher public levels of interest in cannabis as a research topic [9], and increased research accumulation of scientific data have shifted the public perception of the drug from negative to relatively positive viewpoint. Collectively, this has resulted in growth in the number of recreational users and greater volume of legally reportable, real-world data.

Although cannabis is commonly regarded as an anxiolytic, a broader review of the medical literature attests to a range of negative responses to cannabis intoxication. For example, many users experience anxiety and paranoia [10-12], and some studies have also reported an increased risk of developing depression in patients with cannabis chronic use [12-16]. Despite attempts to model causative effects of cannabis towards development of anxiety, stress-vulnerability, and depression, no conclusive evidence has emerged to explain the general observation of increased anxiety disorders among cannabis users [10, 17]. We postulate that variations in cannabis response reside in differences in personality type, particularly that of stress-resilient and stress-vulnerable populations. To address this issue, we used two strains of mice with markedly differing sensitivities to stress and social dominance behavior, which we attribute to representing personality-like features.

The idea of animal personality, temperament or disposition traits has been extensively investigated [18, 19] and is now considered a valid concept in explaining the inter-individual differences in behavioral response. Behavioral responses to stimuli have been demonstrated to be heritable [20] and also consistent throughout the lifetime of indivuals [19]. In the present study, we used selectively-bred mice that represent opposite extremes of the behavioral spectrum of dominance and submissivness and treated them with THC to mimick user exposure. Dominant (Dom) and submissive (Sub) mice simulate different types of animal personalities, with Doms displaying elements of manic-like phenotype and Subs showing depressive-like behaviors [21]. In addition, by using different behavioral paradigms, it was shown that Dom and Sub mice exhibit resilience or sensitivity to stress, respectively [22-25]. The results we present here suggest that personality may dictate the response of individuals to cannabis with regards to development of addictive behaviors.

## 2. Materials and methods

### 2.1 Animal model

Mice were selectively-bred from stock Sabra mice (Harlan Laboratories, Jerusalem, Israel) over 30 generations using a social behavioral paradigm (DSR, see below) which resulted in two animal strains distinct in several measures of social interaction and resource competition: named dominant (Dom) and submissive (Sub) mouse strains [21, 22, 26]. Animals were housed in a colony room (12:12 L:D cycle with lights on 07:00–19:00 hrs, 25±2°C, ambient humidity) in groups of five per cage and provided with standard laboratory chow and water, ad libitum. The experiments were conducted in accordance with NIH/USDA guidelines, under the approval of the Ariel University Institutional Animal Care and Use Committee.

### 2.2 Materials

THC was purified from dried Cannabis flowers based on Wohlfarth et al. 2011 [27]. Briefly, dried flowers were extracted in hexane, filtered, and concentrated under reduced pressure. Flash chromatography was performed on a C18 column using a mobile phase of increasing methanol in water. Fractions containing THC acid were pooled, dried, and decarboxylation performed at 110 C for 1 h. Purity was confirmed using LC-MS.

THC was given in doses of 15 mg/kg or 1.5 mg/kg i.p.

### 2.3 Behavioral experiments

The assessment and treatment scheme for all groups is summarized in Fig. 1. Following verification of Dom or Sub social behaviors (DSR, see below), mice were subjected to conditioned place preference testing (CPP, see below) with THC at either low (1.5 mg/kg) or high (15 mg/kg) doses. Following CPP, mice were allowed a 6-week wash-out period prior to additional behavioral testing (see RS, TST, and FST below). This was done to ensure that mice were not intoxicated during assessments and that all behavioral responses were the result of stable neurological changes. Additional behavioral testing consisted of 3 different stressors applied, one test a day, for 3 consecutive days, starting from lowest to highest stress effect.

**Fig. 1.**
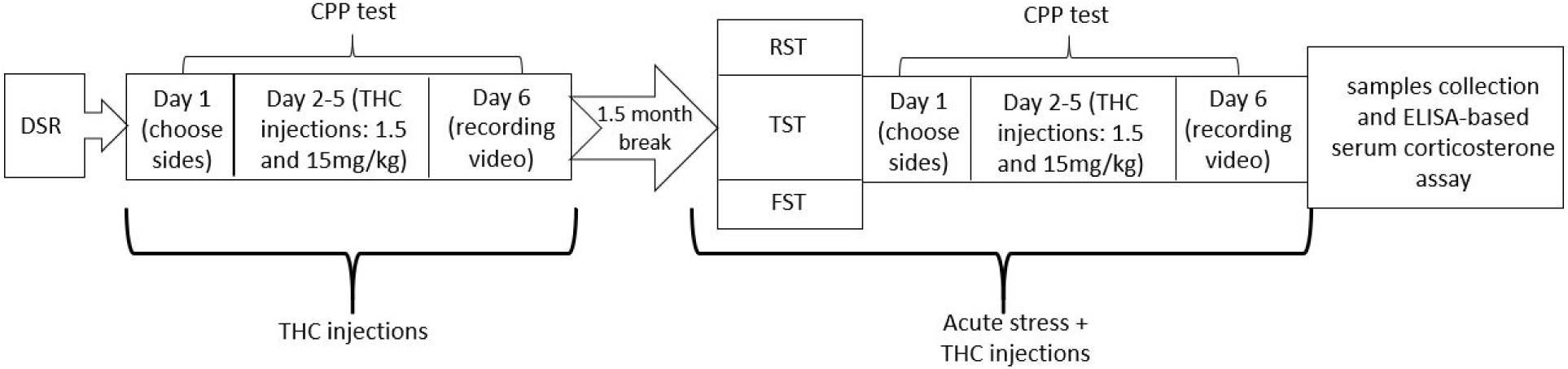
Behavioral assessment and treatment scheme. (RS: restraint stress; TST: tail-suspension test; FST: forced swim test)

#### 2.3.1 Dominant-Submissive Relationship (DSR) Test

The DSR test was used for verifying Dom and Sub strain-specific behaviors as part of selection and colony breeding maintenance. The DSR arena consisted of two identical chambers (l × w × h; 12 × 8.5 × 7 cm) joined by a central, connecting tunnel (27 × 2.5 × 2.5 cm). The competition food target (aqueous solution of 3% milk and 10% sugar) was presented at the tunnel center through a self-refilling well with a small access point to allow feeding by a one mouse in a time only. The end chambers were bordered by removable panels to restrict access to the connecting tunnel and food source until beginning of the test. DSR was conducted for 2 weeks (5 consecutive days/week) with fixed pairs of Dom and Sub mice.

Each testing day, mice were restricted from laboratory chows for 14hr with water provided ad libitum. Dom and Sub mice pairs (6-weeks-old, same gender, each strain) were arranged with individuals of similar weight and placed to the arena for 5 min. Milk drinking time of each animal was manually recorded [21].

#### 2.3.2 Condition Place Preference (CPP)

A conditioned place preference paradigm was used to measure addictive-like behavior [28] by employing an apparatus, which consisted of a plastic box, divided into two compartments (l × w × h, 17 × 15 × 37 cm; one with black and white vertical striped walls and the other with black walls) with a central grey separation section (9 × 15 × 37 cm).

Compartments were separated by removable dividers. On day one, mice were assessed for 20 min without chamber dividers to determine their naïve preference to chamber color and location. On days 2-5 (training days), mice were injected (i.p.) with vehicle solution and then were placed in the closed, preferred outer compartment for 20 min (morning session). Then, mice were injected with THC and were placed in the closed, non-preferred outer compartment for 20 min (afternoon session). Time between sessions was 4hr.

On day 6, (assessment day) mice were placed in the closed central compartment, without dividers, and dwelling time was recorded for each chamber using an EthoVision 3.1 (Noldus, Holland) system.

#### 2.3.3 Restraint Stress (RS)

Mice were exposed to a restraint stress protocol enabling the differentiation of stress effects upon Dom and Sub mice. Mice of each phenotype (Sub, Dom) underwent restraint stress for 1hr using a restriction sleeve, which permitted ease of breathing, but restricted limb movement.

#### 2.3.4 Tail Suspension Test (TST)

The TST is a primary screening test for effects of anti-depressant drugs, which reduce the tail suspension-induced immobility of mice, similar to that observed in the FST. In comparison with the FST, the TST has the advantages of negating the ability of mice to use their natural buoyancy and is considered to be less stress-inducing to the animals [29]. Using a sponge-padded clothespin, animals were suspended by their tails for 6 min from a square stand 30 cm above the table surface. Immobility was recorded manually.

#### 2.3.5 Forced Swim Test (FST)

FST is an acute environmental stressor [22, 30, 31], which measures the time mice spend immobile (non-swim time) following immersion in deep water and is meant to reflect behavioral despair [32, 33]. Mice were placed individually into a glass cylinder (30 × 10 cm) filled 25 cm high with water (25 ± 2°C) for 6 min and immobility time was recorded. Mice were removed from the cylinder at 6 min or earlier if they failed to remain above the water surface. The mice were then dried with paper towels, warmed under a lamp for 10 min, and returned to their home cages. [22, 30].

### 2.4 ELISA-based serum corticosterone assay

Peripheral circulation corticosterone levels were measured in serum samples prepared from trunk blood collected immediately after euthanasia and stored at room temperature for 1hr for erythrocytes clotting and then centrifuged at 10,000 rpm for 10 min, at 4°C. Serum samples were stored at −80°C. Corticosterone concentration was measured using a commercial ELISA kit detecting total serum corticosterone (MS E-5400 LDN, Nordhorn, Germany).

### 2.5 Statistical analysis

For the DSR test, multiple comparisons were performed by two-way ANOVA (column matching with Bonferroni correction for means). For CPP, FST and corticosterone concentrations, multiple comparisons were performed by two-way ANOVA without matching (with Bonferroni correction for means). Means separation test for multiple comparisons were conducted with a Tukey test. Threshold for significance was α=0.05.

## 3. Results

### 3.1 Features of Dom and Sub animals were validated in DSR test

The DSR test was used for selective breeding of mice (>30 generations) to develop animals with strong features of dominance and submissiveness. As depicted in Fig. 2, the DSR test validated behavioral characteristics of selectively-bred Dom and Sub mice.

**Fig. 2.**
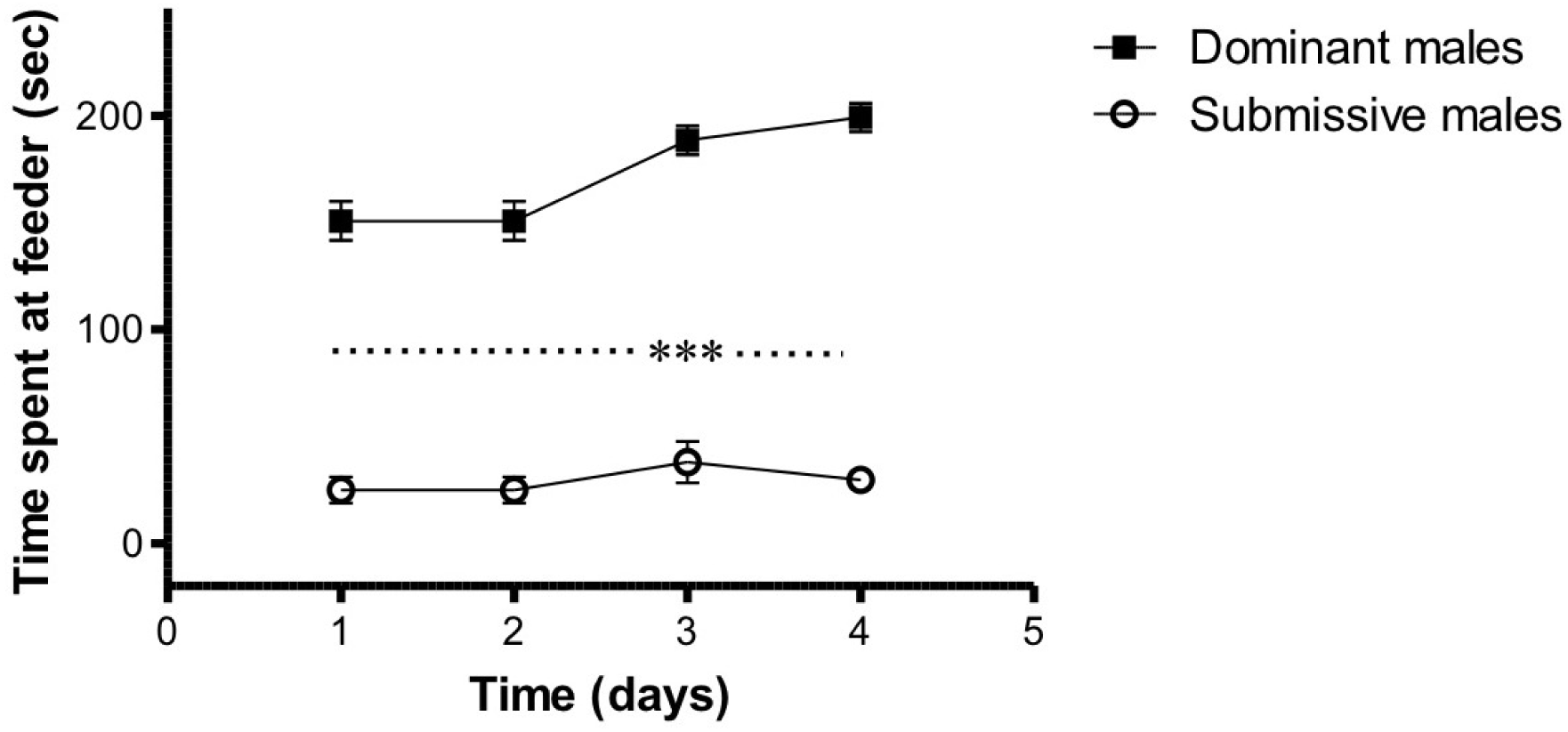
Dominant-Submissive Relationship (DSR) Test. (two-way ANOVA with Bonferroni post hoc analysis, time effect F(1,32)=19.48, P<0.0001; mouse population effect F(1,8) = 265.3, P<0.0001; interaction F(3,24) = 9.738, P<0.0001; n = 5 mice/group), ***P < 0.001, data are presented as delta ± SEM.

### 3.2 Dom and Sub mice differentially responded to THC exposure in the CPP test

According to our hypothesis, personality differences may lead to a differential response to psychoactive compounds. In this experimental design, we assessed the response of socially dominant, stress resilient and socially submissive, stress sensitive animals to THC exposure using the CPP paradigm. The results showed that Sub mice injected with high-dose THC developed strong aversion to the drug. This response was dose-dependent, as low-dose THC did not produce such an effect. In contrast, Dom mice did not exhibit any place preference at any dose of THC injected.

The CPP test was conducted twice, before and after the employment of three consecutive stressors. Surprisingly, acute stress did not influence the development of place preference/aversion, disregarding the doses delivered (see Supplementary Figure A.1). As data obtained from both CPP paradigms didn’t show any statistically significant differences, results were merged (Fig. 3).

**Fig. 3.**
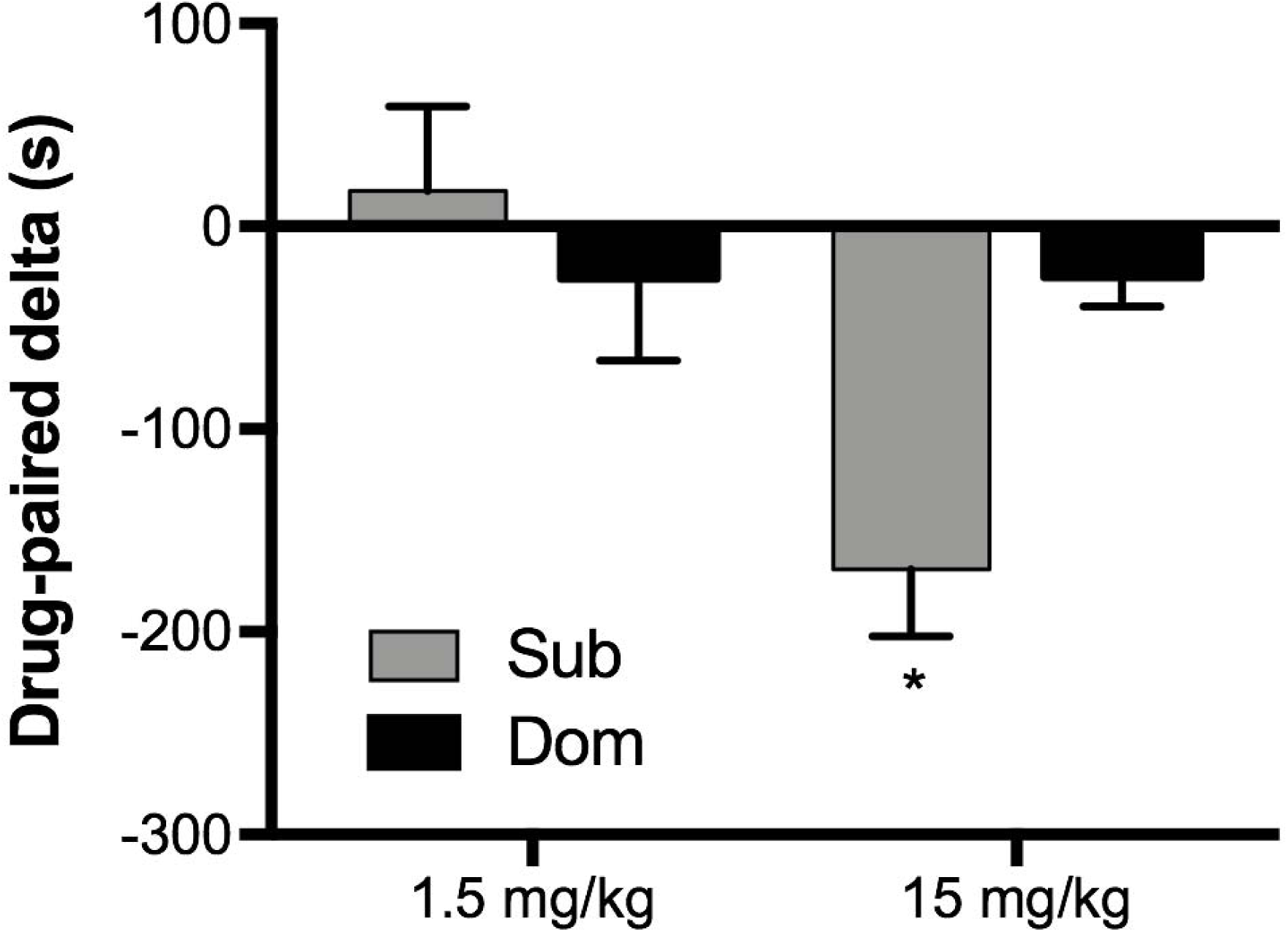
Condition Place Preference (CPP) drug-paired delta time (two-way ANOVA with Tukey’s post hoc analysis, treatment effect F(1,32)=6.801, P<0.05; mouse population effect F(1,32) = 2.001, P = 0.17; interaction F(1,32) = 6.924, P < 0.05; n = 9–10 mice/group), *P < 0.05, data are presented as delta ± SEM.

### 3.3 Exposure to THC differentially altered Dom and Sub mice behavior in the FST test

We anticipate that short-term THC exposure may lead to long-term behavioral effects. Studies show that THC may alter depressive-like behavior [34]; herein, we employed FST to assess whether the effect of THC on depressive-like behavior is personality-dependent. Naïve Sub, but not Dom mice showed high immobility time, which correlates with behavioral characteristics of both populations. Injection of higher THC dose to Dom mice resulted in significant elevation of immobility time, (p<0.001; Fig. 4) suggesting that high THC dose increases depressive-like behavior in socially-dominant individuals.

**Figure 4.**
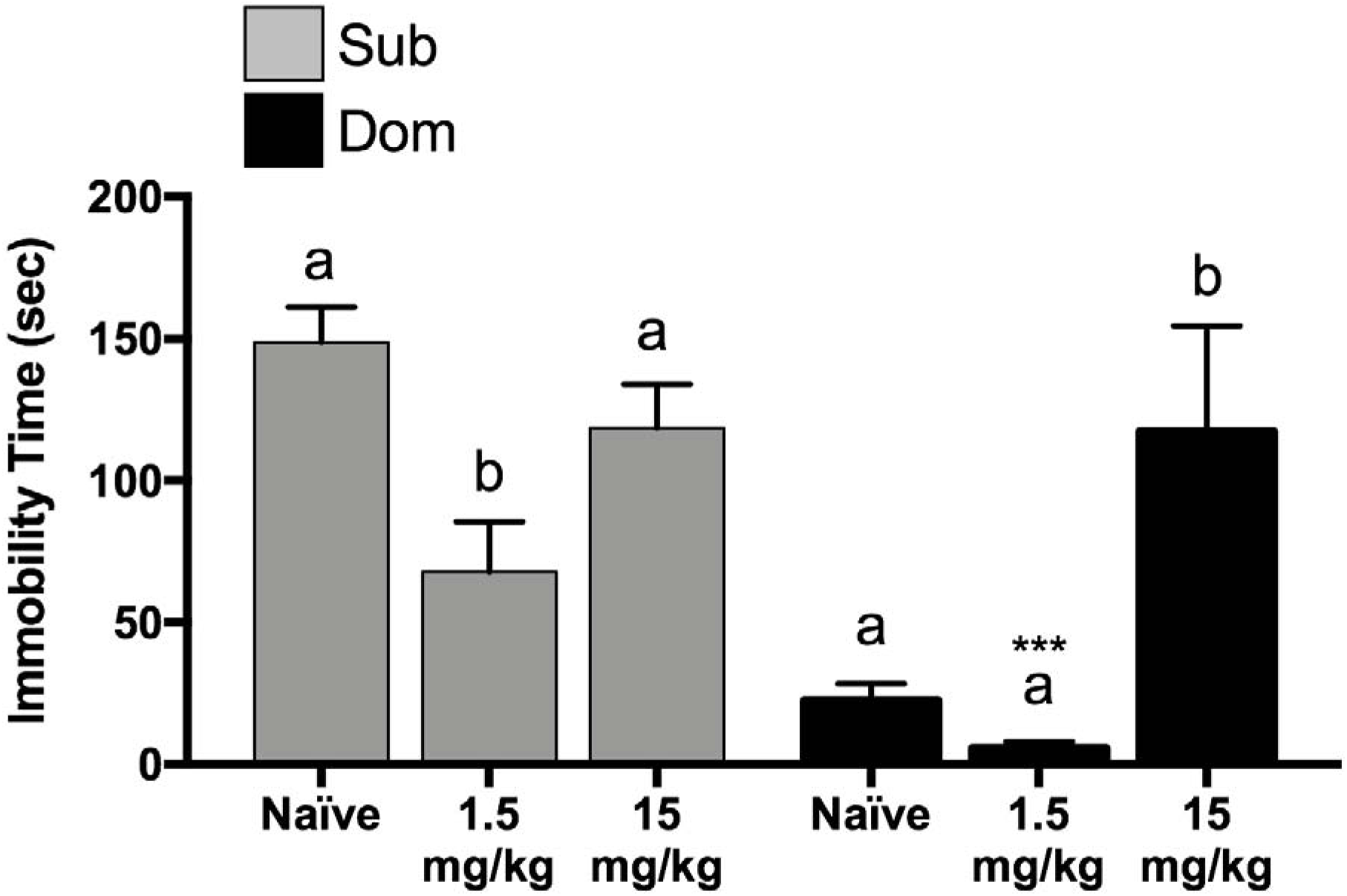
Acute effect of Forced Swim Test (FST) with total immobility time (seconds) in THC-injected mice compared with Naïve groups (two-way ANOVA with Tukey’s post hoc analysis, treatment effect F(2,24)=9.45, P<0.05; mouse population effect F(1,24) = 16.94, P < 0.0 1; interaction F(2,24) = 5.528, P < 0.05; n = 5 mice/group); letters represent results of Tukey means separation test; Naïve Dom vs. 1.5 mg/kg Dom: unpaired t(7)=6.198, ***P < 0.001, data are presented as delta ± SEM.

### 3.4 Exposure to THC dose-dependently altered blood corticosterone concentration

To better understand the observed behavioral changes associated with THC exposure, we evaluated the effect of the drug on the activity of the hypothalamic-pituitary-adrenal (HPA) axis by measuring the concentration of corticosterone (CORT) in serum. CORT concentration was not altered in Dom and Sub mice after injection of low dose THC as compared with naïve mice. However, injection of high dose resulted in significant elevation of CORT concentration both in Dom and Sub groups (Fig. 5).

**Figure 5.**
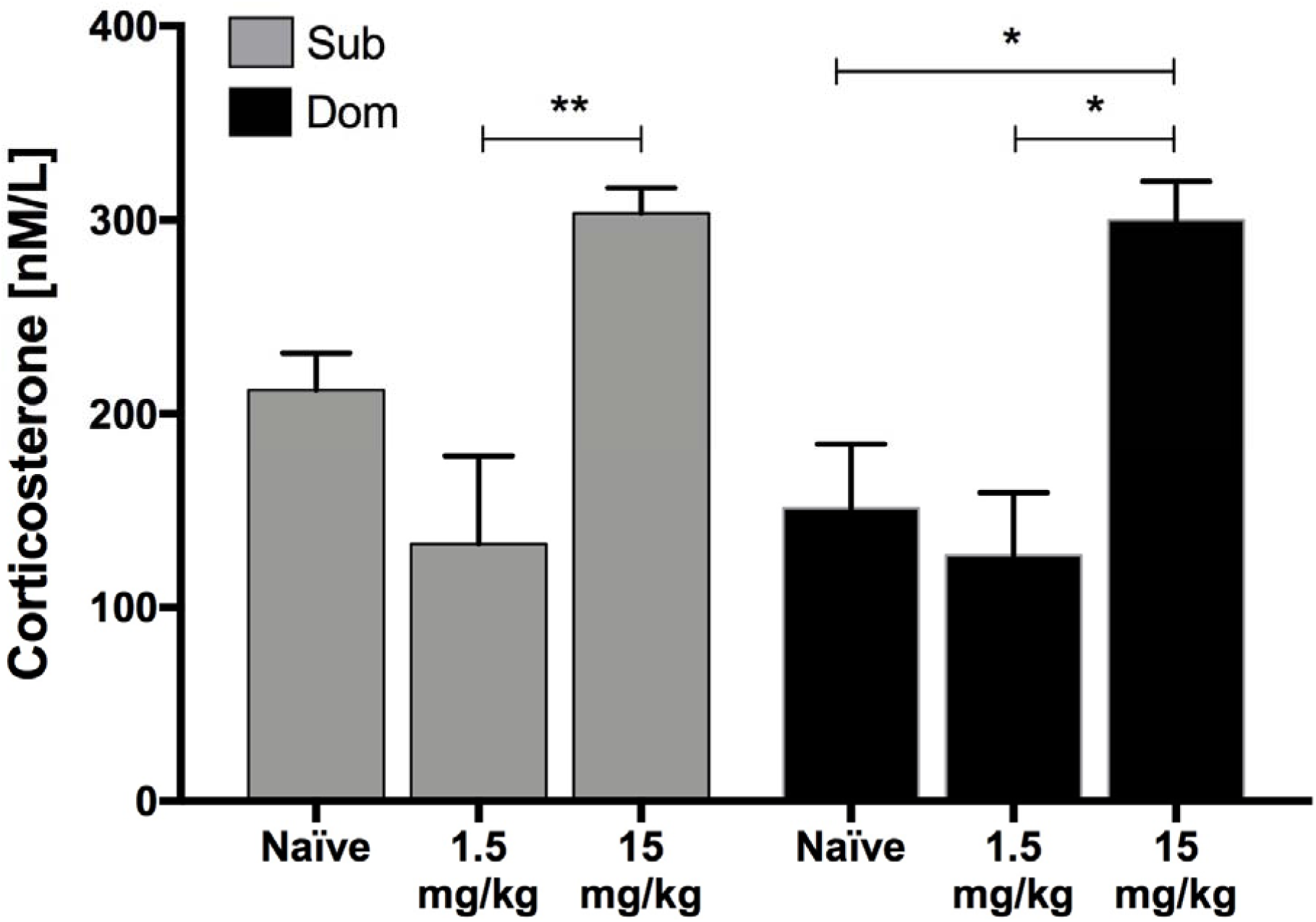
Serum CORT concentration following acute stress and second CPP test (two-way ANOVA with Tukey’s post hoc analysis, treatment effect F(2,24)=18.21, P<0.0001; mouse population effect F(1,24) = 0.97, P=0.33; interaction F(2,24) = 0.6187 P=0.54; n = 5 mice/group; *P<0.05, **P<0.05, data are presented as delta ± SEM.

## 4. Discussion

In this study, we measured long-term outcomes of THC exposure on Dom and Sub mice to better understand the effect of personality on addictive-like behavior. Our previous studies with Dom and Sub mice have established marked differences between strains in response to various stressors [22-25]. Submissive behavior has been linked to increased susceptibility to stress and, presumably, addictive behaviors, whereas resiliency to stress may be associated with dominant personalities [22, 25]. We demonstrated particularly that socially-dominant, stress-resilient and socially-submissive, stress-sensitive mice react differentially to stressogenic factors, antidepressants, and mood stabilizing agents [22, 23, 35, 36]. In addition, these two animal population exhibiting opposite behavioral phenotypes show differing aging-related cognitive impairments and demonstrate significant differences in short- and long-term synaptic plasticity [37]. In currently available studies mice are generally treated with a range of 0.3 to 20 mg/kg THC with 0.3 mg/kg having no effect and 20 mg/kg inducing strong place aversion. Exposure to relatively high doses of THC (5-20 mg/kg ip) results in aversion while low doses (from 1 mg/kg to 4 mg/kg) lead to place preference in most of the studies [38]. In our study we chose 1.5 mg/kg (low) and 15 mg/kg (high) doses of THC as effective and safe [39].

Here, we show that Dom (stress-resilient) and Sub (stress-sensitive) mice react differently following exposure to a high and low dose of THC, implicating the role of social dominance behaviors and/or stress sensitivity in the response to drug exposure.

According to recent studies, chronic use of medical cannabis can lead to various neurological adverse effects depending on the dose of THC and THC-like cannabinoids [40]. The list of neurological symptoms observed after chronic THC exposure is wide and includes seizures, epileptic seizures, headache – the same symptoms that medical cannabis is alleged to cure [40]. Moreover, not only chronic, but acute administration of THC may lead to various psychiatric experiences including anxiety, transient hallucinatory and delusional experiences [41-45]. In one of the first human studies, D’souza et al. (2004) [46] administrated intravenous THC in 2 doses (2.5 mg and 5 mg) to 22 healthy adults in a double-blind, placebo-controlled design. They found that THC induced a psychosis-like experience including symptoms such as: perceptual alterations, anxiety, euphoria, and attention difficulties. In a similar study, Morrison et al. (2009) reported similar effects produced by lower dose of THC [47].

Recent studies also revealed various long-term negative effects of cannabis on mental health including impairment of attention, psychomotor task ability, short-term memory, increased risk of psychoses, depression, and anxiety disorders [48-52]. Our FST results in mice are in agreement with the depression and anxiety aspects observed in human studies. As expected, comparison of naïve mice showed significantly elevated levels of immobility in Sub mice, indicating more prominent depressive-like behavior. This result is consistent with our previous studies and is considered a marker associated with dominant and submissive behavior [22]. We showed here that mouse behavioral patterns were dose-dependently impacted by THC exposure and the effect was long-term and manifested even after a long period of wash-out. Depressive-like behavior was reduced after administration of lower dose of THC in Subs, demonstrating drug stress-relieving and relaxing properties. The lack of significance in Dom naïve vs Dom 1.5 groups can be explained by general very low levels of immobility in Dom naïve group. However, the most prominent response to THC was observed specifically in 15 mg/kg Dom group where the immobility time reached the level of Sub mice, indicating development of depressive-like behavior in Dom individuals despite Dom mice displaying no place preference response to drug as shown in CPP test. These findings are in a good agreement with human studies showing that THC affects individuals differently: some individuals experience relaxation, whereas others develop psychotic states [53, 54].

We postulated that differences in behavioral patterns of Dom and Sub mice after THC injections may reside in their differing sensitivities to stress, therefore we measured the concentration of serum CORT, an indicator of HPA-axis activity. The relationship between elevated serum cortisol and depressive behavior has been established in several studies: Johnson et al. [55] demonstrated that repeated injections of CORT increase depressive-like behavior in rats; in human studies, cortisol concentration was also elevated in response to THC administration, which presumably could result in depressive behaviors in sensitive individuals [56]. In our study, CORT concentration was elevated both in Doms and Subs after exposure to 15 mg/kg THC, yet were unchanged in 1.5 mg/kg group, indicating that higher doses of cannabis may contribute to the development of depressive-like behavior observed in FST. Stimulation of HPA-axis by THC and release of CORT occurs presumably via brain cannabinoid (CB-1) receptors located in the brainstem, namely the locus coeruleus and the nucleus of the solitary tract. Activation of CB-1 receptors by cannabinoids in these regions may modulate noradrenergic activity, resulting in norepinephrine release that has long been known to play a prominent excitatory role in the regulation of the HPA axis. This in turn leads to elevated activity of neurons releasing corticotropin-releasing hormone and, hence, elevated corticotropin concentration. Furthermore, THC may directly activate paraventricular nuclei of the hypothalamus where CB-1 receptors and CRH mRNA are co-expressed [57].

Another experiment conducted in the frame of the current study also revealed personality-based alterations in response to THC. Using the CPP paradigm, we demonstrated that Sub, but not Dom mice, developed strong aversion to THC exposure. CPP tests may demonstrate either place preference or aversion after cannabinoid exposure in test animals, which depends on administered drug dosage [58-62]. Development of aversion to THC in mice may indicate that endogenous cannabinoids are involved in activation of counter-reward pathways that trigger aversion and anxiety. For instance, Bhattacharyya et al. confirmed the anxiogenic role of cannabinoids that was mediated by the modulation of amygdala function through CB-1 receptors [63]. Contrary to our expectations, no effect of acute stress was observed following cannabinoid treatment in both mice groups. One option explaining this phenomenon is that acute stress is not sufficient to affect pathways regulating the development of place preference/aversion in mice after drug exposure.

The finding that the cannabis effect is dependent on individual personality may warrant consideration of cannabis use in relation to medical treatment. We suggest that personality-based behavioral differences should be considered as an essential element of medical cannabis treatment and should be taken into consideration when prescribing and selecting the right dose of THC-containing medications to patients.

## Supporting information

Acute stress did not influence the development of place preference/aversion disregarding the doses delivered (see Supplementary Figure 1)

## 5. Acknowledgements

Research presented herein was conducted in partial fulfillment of the requirements for the doctoral degree of Tetiana Kardash at Ariel University, Ariel, Israel.

## Funding

This research did not receive any specific grant from funding agencies in the public, commercial, or not-for-profit sectors.

## Notes

#### Summary of Updates

Major revision: title changed to better represent the paper's major idea, data reanalyzed using two-way ANOVA, DSR results added

