## Supplementary material for "Link between personality and response to THC exposure": Acute stress did not influence the development of place preference/aversion disregarding the doses delivered (see Supplementary Figure 1)

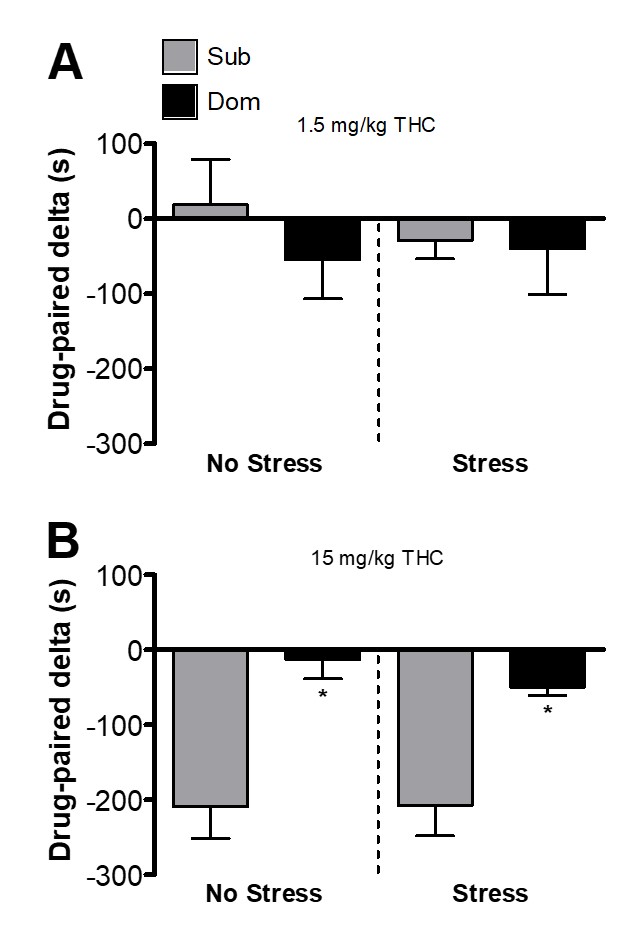

***Appendix 1.*** *Condition Place Preference (CPP) drug-paired delta time (n=5 per treatment).* ***A:*** *No place preference in* *Sub and Dom groups injected with 1.5 mg/kg dose both before (No Stress) and after (Stress) acute stress.* ***B:*** *Sub mice, but not Dom mice, injected with 15 mg/kg dose developed strong aversion to drug both before (No Stress: unpaired t(5)=4.212, *p=0.0084) and after (Stress: unpaired t(5)=3.277, *p=0.0220) acute stress. Data are presented as delta ± SEM.*
